# Genome-wide distribution of 5-hydroxymethyluracil and chromatin accessibility in the *Breviolum minutum* genome

**DOI:** 10.1101/2023.09.18.558303

**Authors:** Georgi K. Marinov, Xinyi Chen, Matthew P. Swaffer, Tingting Xiang, Arthur R. Grossman, William J. Greenleaf

**Affiliations:** Department of Genetics, Stanford University, Stanford, California 94305, USA; Department of Bioengineering, Stanford University, Stanford, California 94305, USA; Department of Biology, Stanford University, Stanford, CA 94305, USA; Department of Plant Biology, Carnegie Institution for Science, Stanford, California 94305, USA; Center for Personal Dynamic Regulomes, Stanford University, Stanford, California 94305, USA; Department of Applied Physics, Stanford University, Stanford, California 94305, USA; Chan Zuckerberg Biohub, San Francisco, California, USA

## Abstract

In dinoflagellates, a unique and extremely divergent genomic and nuclear organization has evolved. The highly unusual features of dinoflagellate nuclei and genomes include permanently condensed liquid crystalline chromosomes, primarily packaged by proteins other than histones, genes organized in very long unidirectional gene arrays, a general absence of transcriptional regulation, high abundance of the otherwise very rare DNA modification 5-hydroxymethyluracil (5-hmU), and many others. While most of these fascinating properties were originally identified in the 1970s and 1980s, they have not yet been investigated using modern genomic tools. In this work, we address some of the outstanding questions regarding dinoflagellate genome organization by mapping the genome-wide distribution of 5-hmU (using both immunoprecipitation-based and basepair-resolution chemical mapping approaches) and of chromatin accessibility in the genome of the Symbiodiniaceae dinoflagellate *Breviolum minutum*. We find that the 5-hmU modification is preferentially enriched over certain classes of repetitive elements, often coincides with the boundaries between gene arrays, and is generally correlated with decreased chromatin accessibility, the latter otherwise being largely uniform along the genome. We discuss the potential roles of 5-hmU in the functional organization of dinoflagellate genomes and its relationship to the transcriptional landscape of gene arrays.

## Introduction

Dinoflagellates are perhaps the most remarkable lineage within the spectrum of known eukaryote diversity, with numerous extreme deviations from the genomic and cellular organization of other eukaryotes, especially regarding their highly unusual nuclei ^1–6^. They are also a very diverse, successful, and ecologically important group that includes numerous photosynthetic lineages, free living heterotrophs, and even parasites, playing a major ecological role in marine ecosystems. The best known such example is the endosymbiotic association of Symbiodiniaceae dinoflagellates ^7^ with reef-building corals. The photosynthetic capability of the dinoflagellate symbionts provides the metabolic foundation for the highly biologically diverse reef ecosystems ^8^, and the expulsion of these symbionts from their host cells upon heat stress causes coral “bleaching” and the eventual death of coral reefs ^9^, an increasingly acute problem in the modern world due to the effects of global climate change ^10^.

The list of unorthodox features of dinoflagellate nuclei is long ^1,3,4^. Dinoflagellate chromosomes exist in a permanently condensed liquid crystalline state throughout most of the cell cycle, and are characterized by an unusually low protein-to-DNA ratio (1:10, compared to 1:1 in other eukaryotes ^2,15^). This condensed, protein-poor structure is caused by the loss of nucleosomal histones as the main packaging component of chromatin. This role has instead been taken over by a distinct set of proteins – small dinoflagellate-specific virus-derived nucleoproteins (DVNP) and histone-like proteins (HLPs) ^16–24^ – that appear to have been acquired through horizontal gene transfer from viruses and bacteria, respectively ^25,26^. Such chromatin composition is an extreme departure from the norm for an eukaryote, as nucleosomal chromatin is otherwise universal ^27^. Furthermore, the four core histones – H2A, H2B, H3 and H4 – are the most conserved of all eukaryotic proteins, a result due not just to their role in packaging DNA, but also because they, especially in their N-terminal tails, are subject to extensive posttranslational modifications (PTMs) that serve as the basis of the so-called “histone code” ^28^. This code plays important roles in all chromatin-related biological processes, such as transcription, gene regulation, replication, DNA repair and others. Dinoflagellates have not lost histones – in fact multiple and highly diverse histone genes are retained in all dinoflagellates for which genomic data is available ^29^ – but these histone proteins are highly divergent from those of other eukaryotes and such divergence makes it unclear what role these proteins might play in dinoflagellate nuclei. DVNPs and HLPs appear to have been acquired at different points in dinoflagellate evolution ^26^ – DVNPs are found in all dinoflagellates, but HLPs are absent in early branching lineages and common only to core dinoflagellates. Another curious feature of dinoflagellate nuclei is the high abundance of divalent cations such as Ca^2+^ and Mg^2+^, which have been proposed to play a role in the stabilization of chromosome structure ^30–33^.

Genome organization in dinoflagellates also represents a highly derived state, as their genes are organized into long unidirectional gene arrays ^34–37^, presumably transcribed as a single unit. *Trans*-splicing is ubiquitous in the group, with mature mRNAs generated through the addition of a spliced leader (SL) sequence ^34,38,39^. Transcriptional regulation is thought to be largely absent, with all genes transcribed at all times. The primary mode of gene regulation is presumed to be at the level of translation and/or RNA stability.

We still know very little about the inner workings of these remarkable nuclei, and most of the numerous fascinating questions regarding how eukaryotic transcription, replication and DNA repair occur and are regulated in the absence of histones and within permanently condensed liquid crystalline chromosomes remain unanswered. Recently, we and others ^40,41^ began to unravel some of these mysteries by applying three-dimensional genome conformation mapping using Hi-C ^42^ to the members of Symbiodiniaceae *Breviolum minutum* and *Symbiodinium microadriaticum*, showing that the genome is folded into distinct topologically associating domains coinciding with pairs of divergent gene arrays and separated by the points where convergent gene arrays meet (termed “dinoTADs”). These domains appear to be the product of strong transcription-induced supercoiling in a context of extremely long transcriptional units and the absence of histones.

Earlier, de Mendoza et al. ^43^ mapped the distribution of the frequently found in eukaryotes 5-mehylcytosine (5mC) modification in two members of the Symbiodiniaceae – *Fugacium kawagutii* and *Breviolum (Symbiodinium) minutum*. An unusual compared to other eukaryotes pattern of uniform hypermethylation throughout the genome was observed.

Two other still unanswered questions concern the potential roles and genomic distribution of the 5-hmU modification in dinoflagellate genomes and the characteristics of the chromatin accessibility landscape. While it appears that nucleosomes are not the main packaging component of dinoflagellate chromatin, it is not known whether DVNPs and/or HLPs might provide similar levels of physical protection of DNA. Past studies have suggested that DVNPs bind to DNA with similar affinity to histones ^25^ but HLPs, although they can compact DNA in a concentration-dependent manner, have weaker affinity than histones ^44–46^. However, whether and how these proteins confer distinct chromatin states through their association with DNA is not known. Thus, whether dinoflagellate genomes are uniformly accessible given the lack of histones, whether distinct open chromatin region exist as in conventional eukaryotes, and whether perhaps the inverse phenomenon is observed – localized areas of decreased accessibility – is an open question.

That dinoflagellates contain the otherwise highly unusual for eukaryotes 5-hmU modification (present in abundance and discovered originally in some phages ^47^) was first realized in the 1970s ^48–50^. It was found that unexpectedly large fractions of thymines (T) in the genome of various species are replaced by 5-hmU – 12% in *Exuviaella cassubica* ^50^, 12% in *Symbiodinium microadriaticum* ^50^, 37-38% in *Crypthecodinium cohnii* ^50,51^, 62% in *Amphidinium carterae* ^50^, 62.8% in *Prorocentrum micans* ^52^, and 68% in *Peridinium triquetrum* ^50^. What functions 5-hmU might have is not known, but it has been suggested that it enhances the flexibility and hydrophilicity of double-stranded DNA ^53^, especially in some sequence contexts ^54–56^.

Curiously, 5-hmU has also been observed in non-negligible quantities ^57^ in another major lineage of eukaryotes – the important parasitic clade Kinetoplastida. There are many parallels between the genomic organization of dinoflagellates and kinetoplastids ^58^ – although kinetoplastids have conventional nucleosomal chromatin, they too have lost transcriptional regulation as a primary mechanism for controlling gene expression and their genes are also organized into long arrays ^59–62^, with mature mRNAs being the product of *trans*-splicing ^63–67^. These properties are shared with other members of the larger Euglenozoa lineage that have been studied, such as *Euglena gracilis* ^68^. However, in kinetoplastids 5-hmU appears to be simply a precursor to the synthesis of the larger modification *β*-D-Glucopyranosyloxymethyluracil, better known as base J ^69–71^, which does play a significant role in their genomes. Base J replaces about 1% of thymines and is predominantly found in repetitive DNA, especially in telomeric regions ^72–74^, but more importantly, it also demarcates the boundaries between gene arrays ^75^ and likely prevents transcriptional readthrough events ^76,77^. The free living *Euglena* also has base J ^68^, and thus 5-hmU too. It is likely that so do all members of the larger Euglenozoa grouping (comprising the kinetoplastids, euglenids, diplonemids and a few smaller clades), but that has not yet been directly demonstrated.

To answer these questions, we mapped the 5-hmU distribution in the genome of *B. minutum*, finding that it is enriched over certain repetitive element classes and often around the boundaries between gene arrays. In contrast, chromatin accessibility is anti-correlated with elevated 5-hmU levels; this inverse relationship is specifically strong around gene array/dinoTAD boundaries, pointing to potential localization of histones (or other proteins that protect DNA) to regions enriched for 5-hmU (and thus conferring them greater protection from transposase insertion). We do not detect increased accessibility associated with transcription start sites (TSSs), and generally we do not observe strongly localized DNA accessibility peaks in the genome comparable to those in metazoans. These results provide a foundation for the future detailed understanding of the organization of transcription in dinoflagellates and its interplay with DNA modifications.

## Results

### Mapping 5-hmU and chromatin accessibility in *B. minutum*

In order the map the distribution of 5-hmU in the *B. minutum* genome, we first adapted the MeDIP (Methylated DNA ImmunoPrecipitation) protocol ^78^ for mapping DNA methylation using high-throughput sequencing (MeDIP-seq ^79^). We used an antibody specific to 5-hmU (see Methods) and a spike-in control to confirm specific enrichment of 5-hydoxymethyluracil. As mammalian genomes do not contain appreciable amounts of 5-hmU, we used a mixture of human and *B. minutum* genomic DNA (gDNA) as input to the MeDIP procedure, and we also sequenced three different controls – input DNA, “depleted” DNA (the supernatant remaining after the immunoprecipitation step), and an IgG control (using only beads with no primary antibody). We observed that the fraction of human reads decreased *∼*2*×* after 5-hmU MeDIP relative to controls (Figure 1A), confirming the specific enrichment of dinoflagellate DNA. We also made an interesting observation – 5-hmU MeDIP is also *∼*2*×* enriched for multimapping reads compared to the controls (Figure 1B), hinting that 5-hmU might be preferentially associated with repetitive elements.

**Figure 1:**
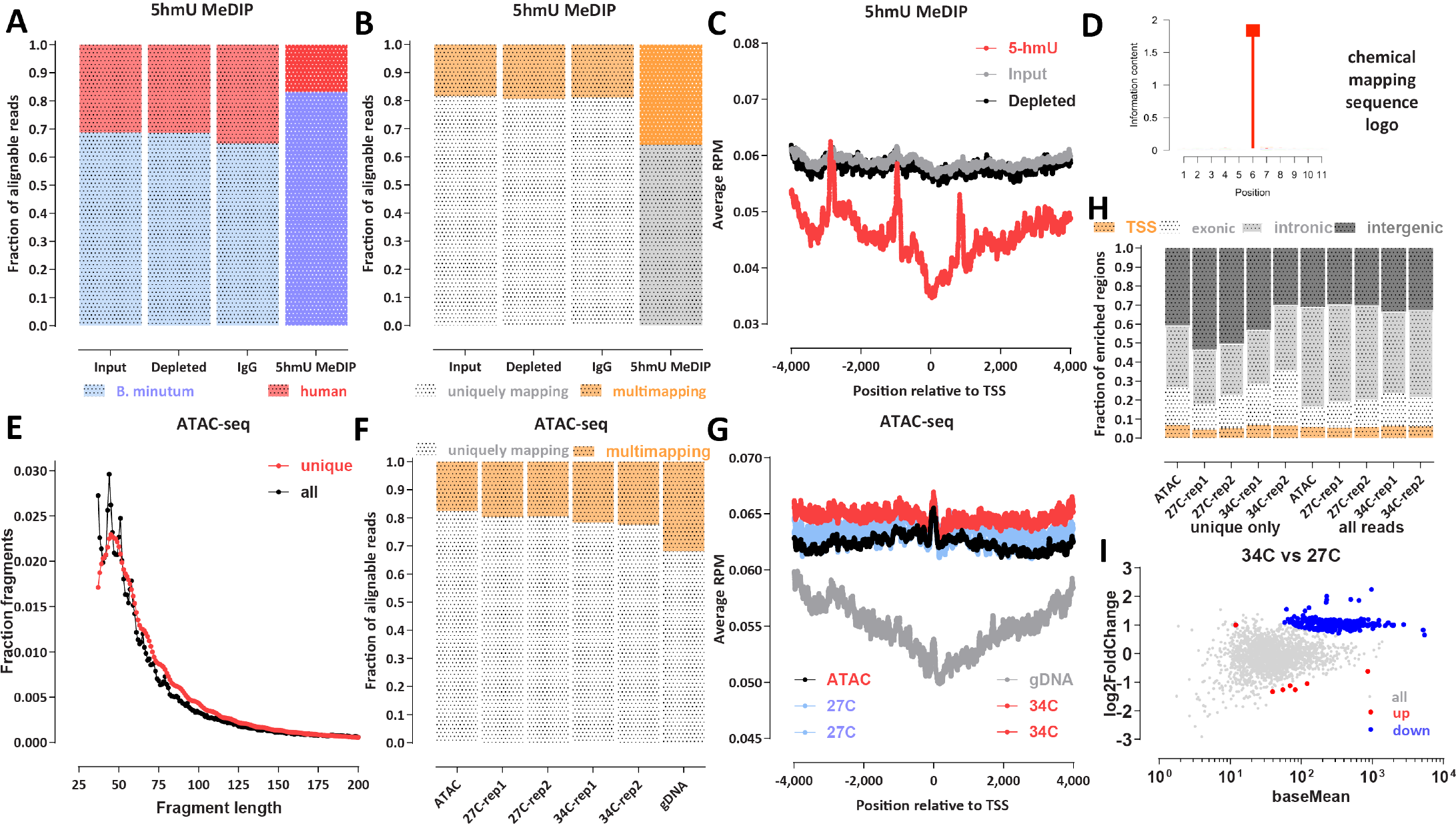
Mapping the 5-hmU and chromatin accessibility landscape in *B. minutum*. (A) Proportions of human and *B. minutum* gDNA in 5-hmU Methylated DNA immunoprecipitation sequencing (MeDIP-seq) and control libraries. A mixture of human and dinoflagellate gDNA was used as input to MeDIP-seq experiments, and the fraction of reads that map to each genome is shown. The 5-hmU MeDIP-seq library is enriched for dinoflagellate reads confirming the specificity of 5-hmU pull down. (B) Proportion of multimapping reads in 5-hmU MeDIP-seq and control libraries. The 5-hmU MeDIP-seq library exhibits a higher fraction of multimapping reads, suggesting that 5-hmU is enriched over repetitive elements. (C) Metaprofiles of 5-hmU and control libraries signal over *B. minutum* transcription start sites/gene starts. (D) Basepair-resolution chemical mapping of 5-hmU does not reveal a sequence motif associated with the modification in *B. minutum*. (E) Fragment length distribution of *B. minutum* ATAC-seq datasets. Shown are uniquely mapping reads alone as well as all reads that can be mapped. (F) Proportion of multimapping reads in *B. minutum* ATAC-seq datasets as well as a control genomic DNA (gDNA) library. (G) Metaprofiles of ATAC-seq signal over *B. minutum* transcription start sites/gene starts as well as the gDNA control. (H) Distribution of ATAC-seq regions of enrichment relative to annotated genomic features. (I) Differential accessibility analysis for the 27 ^*°*^C and 34 ^*°*^C conditions.

We did not observe enrichment or 5-hmU around the starting positions of genes (Figure 1C) – in fact we observe a slight depletion *±*1-kb around gene starts (note that the three spikes observed in the plot are an artefactual result due to the presence of collapsed repeats in the current *B. minutum* assembly; see further discussion on this topic below).

We also deployed an orthogonal method for mapping 5-hmU at base-pair resolution using chemical conversion of 5-hmU into cytosine C (see the Methods section for details). In contrast to the 5mC modification in mammals, which is found specifically in a CpG context, we do not find any sequence preference for T bases modified into 5-hmU in *B. minutum* (Figure 1D). We note that early studies from the 1980s reported that 5-hmU preferentially replaces thymines in TA and TC sequence contexts ^51^, and we do not recover such a preference in our datasets; it is possible that such preferences indeed exist in *Crypthecodinium cohnii*, which was assayed by those studies, but not in Symbiodiniaceae, or that the discrepancy is due to methodological differences. Because the current chemical mapping protocol does not provide for 100% conversion rate of 5-hmU modified bases it is not possible to estimate the absolute levels of 5-hmU in the *B. minutum* genome based on chemical mapping data alone.

To map the *B. minitum* chromatin accessibility land-scape, we utilized ATAC-seq ^80^ (Assay of Transposase-Accessible Chromatin using sequencing), specifically in its omniATAC ^81^ modification (see Methods). We generated a very deeply sequenced (*∼*130 million mapped reads) library from actively growing cells as well as two replicates each for cells grown at the usual temperature of 27 ^*°*^C and heat-stressed cells that were incubated at 34 ^*°*^C.

In eukaryotes with nucleosomal chromatin, ATAC-seq libraries sequenced in a paired-end format display a characteristic nucleosomal signature in their fragment length distribution, with a subnucleosomal peak at *≤∼*120 bp, a prominent mononucleosomal peak, and a weaker dinucleosomal peak. *B. minutum* ATAC-seq only displays a peak at short fragment lengths (*∼*60 bp), with no nucleosomal peaks (Figure 1E). Thus, we conclude that wherever they are found in the genome, nucleosomes apparently are of too low abundance to substantially affect the overall fragment length distribution, while DVNPs and HLPs do not form structures consisting of multiple closely positioned proteins that strongly protect against transposition. We also observe a modest depletion of multimapping reads in ATAC-seq libraries relative to a matched naked gDNA control (Figure 1F), i.e. the opposite trend of that observed for MeDIP-seq. ATAC-seq signal is also not enriched around gene start positions (Figure 1G).

Genome browser inspection of ATAC-seq and gDNA controls (Supplementary Figures 1 and 2) revealed that the available *B. minutum* assembly includes multiple collapsed repeats, i.e. regions that exist in multiple copies in the actual genome but are only present in the assembly as a single copy (or as many fewer copies than their actual abundance in the genome). This complicates the interpretation of sequencing datasets as these regions appear as artificial “peaks” if analysis is not carried out against a proper control. Therefore, we performed all subsequent analysis as a comparison against matched input or negative gDNA controls. The regions of enrichment over gDNA that we identified did not show a concentration around gene starts/TSSs (Figure 1H)., and they show overall lower enrichment over background/controls than ATAC-seq peaks in human datasets ^82^ (Supplementary Figure 3), i.e. we do not really observe strongly localized chromatin accessibility as in other eukaryote genomes. Comparing the heat stressed (34 ^*°*^C) and normal temperature (34 ^*°*^C) conditions did not reveal large scale changes in the chromatin accessibility landscape (Figure 1I).

### Heterologous expression of DVNPs has a modest effect on chromatin organization in the yeast *S. cerevisiae*

Previous studies had examined the effect of DVNPs on chromatin structure by expressing a DVNP (*Hematodinium sp*. DVNP.5) in the yeast *Saccharomyces cerevisiae* ^83^. The resulting changes in the chromatin landscape (measured using MNase-seq) were reported to reveal nucleosome disruption, while overall the expression of the DVNPs had a negative effect on cell growth, likely because it impaired transcription. We sought to replicate and expand on these results by expressing several DVNPs in *S. cerevisiae* and carrying out ATAC-seq as well as single-molecule footprinting (SMF ^84,85^; providing information about the absolute levels of accessibility/protection along the genome associated with DVNP occupancy).

We hetorologously expressed (see Methods) three different DVNPs in yeast – the previously assayed *Hematodinium sp*. DVNP.5 as well as *Hematodinium sp*. DVNP.12 and *B. minutum* DVNP symbB.v1.2.006931. *Hematodinium sp*. DVNP.5 had a negative effect on growth, as previously reported, but much more strongly so than the other two. We carried out ATAC-seq and SMF using an internal control in all experiments – *Candida glabrata* cells, which we used to account for experimental variation, as previously described ^86^.

ATAC-seq did not reveal dramatic changes in the accessibility landscape upon DVNP expression (Supplementary Figures 5 and 4) except perhaps for a slight decrease in the height of some peaks. On the other hand, SMF data showed a decrease in accessibility around TSSs and reduced strength of nucleosome positioning (Supplementary Figure 5), broadly consistent with the previous MNase-seq results suggesting that nucleosome disruption is induced by DVNPs ^83^. This disruption is, however, apparently not sufficient to dramatically reshape the accessibility landscape and represents a moderate quantitative rather than a major qualitative alteration.

### Inverse correlation between 5-hmU and chromatin accessibility

Next, we examined the distribution of 5-hmU and chromatin accessibility around other available genomic features. We noticed that in many cases (although this is not an exclusive association) 5-hmU is enriched around the boundaries of dinoTADs while ATAC-seq shows decreased accessibility in those same regions (Figure 2A). We generalized this observation by evaluating the global ATAC-seq distribution around dinoTAD boundaries (Figure 2C-E) and found that indeed ATAC-seq is globally depleted nearby these locations, while MeDIP-seq is enriched and 5-hmU chemical conversion rate is also elevated. The observed anti-correlation between chromatin accessibility and 5-hmU is specifically strong around dinoTAD boundaries while we do not observe substantial inverse correlation between the two in the middle of dinoTAD domains (Figure 2H). However, these observations are trends and not universal patterns, as a number of gene array boundaries do not show strong MeDIP enrichment and ATAC-seq depletion.

**Figure 2:**
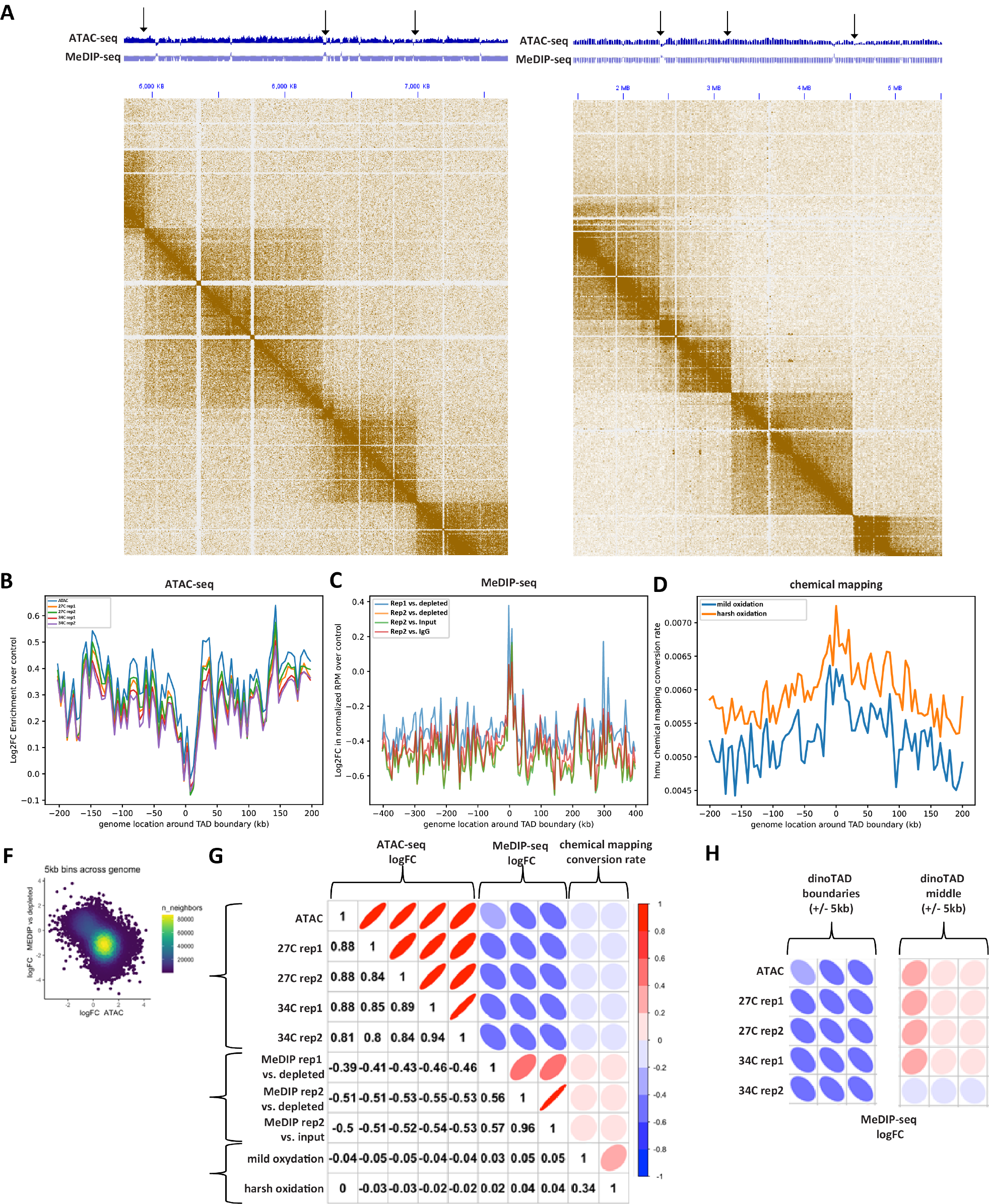
Inverse correlation between 5-hmU and chromatin accessibility and association with dinoTADs boundaries in the *B. minutum* genome. (A-B) Representative snapshots of the distribution of 5-hmU enrichment and decreased chromatin accessibility relative to dinoTAD boundaries. (C) Depletion of ATAC-seq signal around dinoTAD boundaries. (D) Enrichment of MeDIP-seq signal around dinoTAD boundaries. (E) Increased 5-hmU chemical mapping conversion rate around dinoTAD boundaries. (F-G) ATAC-seq and MeDIP-seq are generally anti-correlated (calculated for 5-kbp bins over the whole genome) (H) ATAC-seq and MeDIP-seq are specifically strongly anti-correlated around dinoTAD boundaries.

### Association of 5-hmU and chromatin accessibility with repetitive elements

Because of the previously noted enrichment and depletion of multimapping reads in 5-hmU and ATAC libraries, respectively, we next aimed to identify which, if any, repetitive elements might be specifically associated with 5-hmU and/or ATAC. We first examined the distribution of annotated repetitive elements (see Methods for details) around dinoTAD boundaries (Figure 3A-C), and found no specific preference at dinoTAD boundaries neither for repeats as a whole, nor for any specific repeat family. However, Maverick DNA elements did exhibit strong enrichment around the edges of dinoTADs (Figure 3C). Maverick elements are also known as Polintons, are typically 15-40 kbp in size, and often encode putative viral capsid proteins, suggesting that they might form virions under some conditions ^87–91^, a view supported by the large abundance of Polinton-like viruses reported in aquatic ecosystems ^92^. Maverick does not account for all dinoTAD boundaries though – while a majority of TAD boundaries show 5-hmU enrichment, Maverick elements are found around only *∼*11% of them (Supplementary Figure 6).

**Figure 3:**
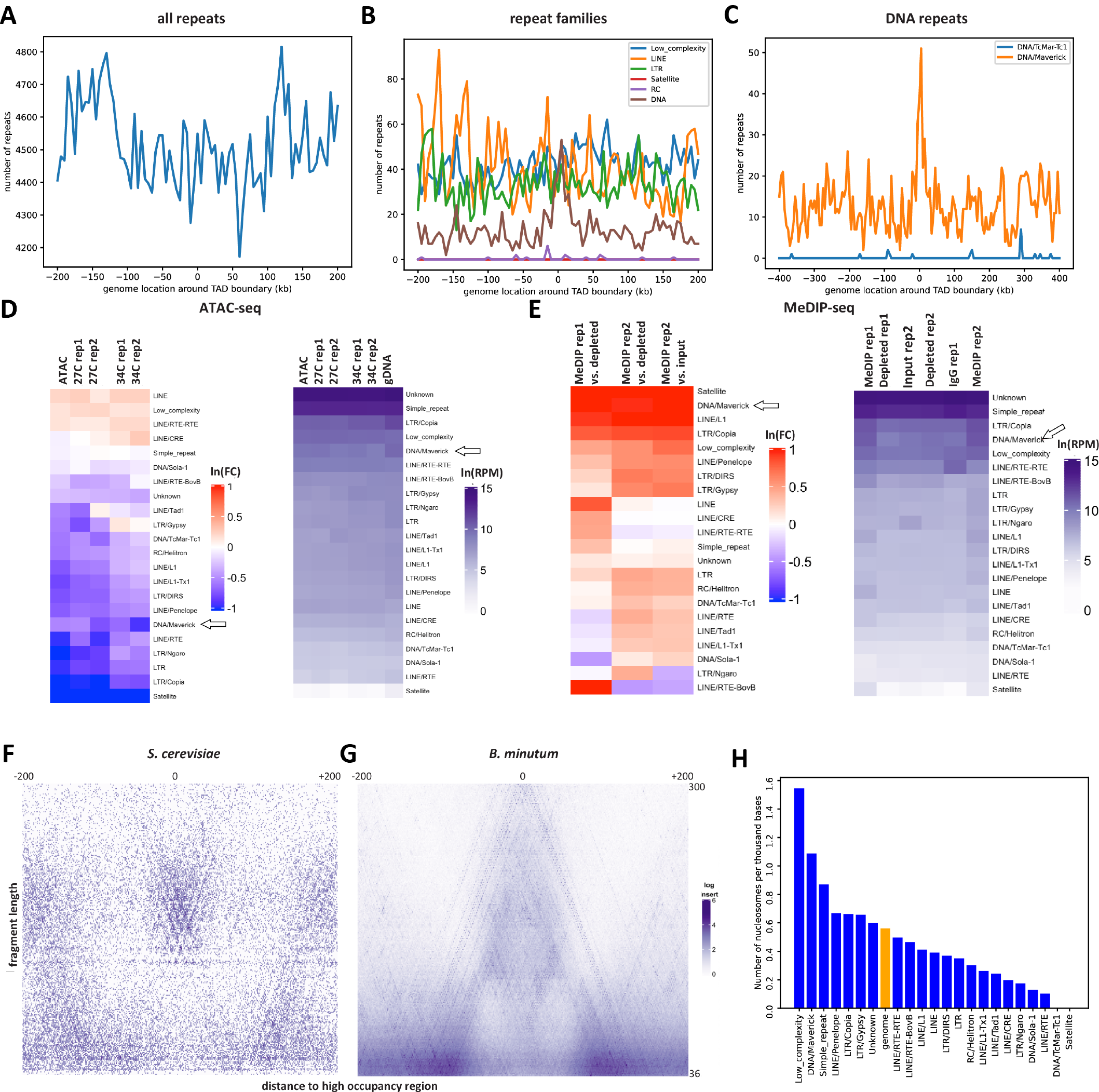
Association of 5-hmU and chromatin accessibility with repetitive elements in the *B. minutum* genome. (A-C) Distribution of all repeats, individual repeat families, and some DNA elements around dinoTAD boundaries. (D) ATAC-seq enrichment/depletion over repetitive elements. (E) MeDIP-seq enrichment/depletion over repetitive elements. (F-G) V-plot ^94^ around positioned nucleosomes in S. cerevisiae (for comparison) and *de novo* identified putative positioned nucleosomes in *B. minutum*. (H) Enrichment/depletion of positioned nucleosomes over repetitive elements.

Global analysis of ATAC-seq depletion/enrichment over repetitive elements (Figure 3D) showed that most repeats are depleted for accessibility, with Copia LTR and Maverick DNA elements most highly abundant in the gDNA control relative to ATAC-seq sample. The exceptions are CRE and RTE-RTE LINE elements. MeDIP-seq data reveals a generally inverse picture – most repeats are enriched in the MeDIP libraries, apart from some LINE elements (Figure 3E), with Maverick/Polinton DNA repeats most strongly enriched for the 5-hmU modification.

These results point to increased protein occupancy and elevated 5-hmU levels over repetitive elements such as Mav-erick. We therefore asked whether we can specifically find nucleosomes corresponding to certain repeat classes. We utilized the nucleoATAC algorithm ^93^ to identify positioned nucleosomes genome-wide in the *B. minutum* genome (see Methods; with the caveat that the nucleoATAC algorithm was designed to look for eukaryote-like nucleosomes and we do not know if this is still the case in dinoflagellates). We identified 30,107 low-resolution and 2,166 high-resolution putative positioned nucleosomes; these are overall not preferentially located to well defined general genomic features such as dinoTAD boundaries (Supplementary Figure 7). V-plot analysis ^94^ of the fragment distribution around positioned nucleosomes revealed an A-shaped structure, with a peak in the 120-160 bp range (this fragment length is higher for the smaller set of high-resolution nucleosomes; Supplementary Figure 7), flanked by very short fragments. This observation is distinct from what is observed in other eukaryotes such as yeast (Figure 3F-G), where multiple nearby nucleosomes are visible. We interpret these structures as arising from a single positioned protective feature, quite possibly a histone-based nucleosome, without other strongly positioned nearby nucleosomes. We note that these observations are not explainable by mappability biases (i.e. only a single nucleosome is observed because all adjacent sequences are not mappable), as we carried out this analysis while allowing for multimapping reads and the center point of the putative positioned nucleosomes is in fact slightly less uniquely mappable than the flanks (Supplementary Figure 8).

Strikingly, Maverick DNA elements are preferentially enriched for positioned nucleosomes, at *∼*2*×* the genomic average (Figure 3H). This observation corroborates the depletion of ATAC-seq signal observed over these elements.

## Discussion

In this study, we provide the first global maps of the distribution of the 5-hmU modification and chromatin accessibility in a dinoflagellate species (*B. minutum* in the Symbiodiniaceae clade). Our results point to the preferential enrichment for 5-hmU over certain classes of repetitive elements and also around the boundaries of the previously identified dinoTAD topologically associating domains that also coincide with the points of convergence of the long unidirectional arrays into which dinoflagellate genes are organized. In contrast, chromatin accessibility is depleted in those same areas and is generally anti-correlated with high levels of 5-hmU. We do not observe strong accessibility peaks as seen in eukaryotes with conventional nucleosomal chromatin, nor do we see any preferential accessibility around transcription start sites, suggesting that most of the dinoflagellate genome is not protected by strings of nucleosomes and is generally uniformly physically accessible. We also do not see evidence for large-scale domains of preferential accessibility/protection; previously ATAC-see ^95^ was used to visualize through microscopy the accessi-ble chromatin in the cells of the basal dinoflagellate *Hematodinium* ^13^, which suggested the existence of large-scale “open” and “closed” compartments, perhaps encompassing whole chromosomes. However, no ATAC sequencing data is available for *Hematodinium*, thus we do not know how these open/closed compartments translate into genome-wide sequencing profiles. It is possible that the open compartment is fully open and the closed one is fully closed, i.e. without distinct loci of open chromatin, resulting in a similar average genome-wide profile to what we observe. Alternatively, chromatin structure in basal and core dinoflag-ellates (Symiodiniaceae belonging to the latter) may differ substantially. The application of single-molecule foot-printing techniques to dinoflagellates should be able to resolve these possibilities in the future ^85,96^ though at present it is hampered by the high sequencing coverage over the very large dinoflagellate genomes that is required. We do identify several thousand putative positioned nucleosomes; however, these, if they are confirmed to be indeed histone-based nucleosomes, appear to be isolated and not parts of larger-order structures. An interesting trend that emerges is the association of elevated 5-hmU, decreased chromatin accessibility and increased frequency of positioned nucleosomes over certain repetitive elements, in particular Maverick/Polinton DNA elements, which are also enriched over dinoTAD boundaries. This is, however, by no means an absolute rule as not all dinoTAD boundaries are associated with such features.

Nevertheless, it is tempting to draw parallels between these initial observations in dinoflagellates and what is known in much more details for kinetoplastids ^58^. The latter group shares the same general loss of transcriptional regulation as a primary mechanism for modulating gene regulation and the organization of the genome into long unidirectional gene arrays, and in the kinetoplastids base J demarcates the regions between these arrays. Base J is synthesized through 5-hmU as an intermediate, and thus 5-hmU also is localized to the same regions of the genome in the cases where it has been measured (e.g. in *Leishmania* ^97,98^). It is possible that it may play an analogous role in dinoflagellates, even though they have not evolved the further chemical elaboration needed that for base J synthesis.

However, such a speculation still leaves many unanswered questions. What is the mechanistic role of 5-hmU? In our previous work in which dinoTAD structures were discovered ^40^, we showed them to depend on transcriptional activity and to disappear upon blocking transcription, i.e. they are most likely the product of extreme transcription induced DNA supercoiling. At the same time, 5-hmU has been reported to increase the flexibility of the DNA double helix ^53–56^, which may suggest a possible role for 5-hmU in alleviating the supercoiling stress under which dinoflagellate genomes appear to exist. However, the mechanistic details of such a link are currently unclear.

Why does 5-hmU vary so much between different dinoflagellate species – from 12% to 68% where it has been assayed – and where is it located in the genome in the extreme cases? The preferential localization to dinoTAD boundaries suggested from our work is consistent with genome-wide rates of 5-hmU on the lower end of this spectrum (which is also what the available data for other Symbiodiniaceae points to ^50^), as array boundaries are fairly localized and encompass a minor fraction of the whole genome. However, the *B. minutum* genome is relatively small for a dinoflagellate – only on the order of 1 Gbp – while other species have much larger and more repeat-rich genomes, which might be related to higher overall 5-hmU levels given the preference of 5-hmU for repetitive elements that we observe. To better understand its properties and functions, it will be important to assay 5-hmU in a wide variety of dinoflagellate species with diverse genomic characteristics.

Similarly, it will also be vital to obtain very high-quality genome assemblies to work with. For example, in our current work we have not been able to test whether 5-hmU is strongly associated with telomeres the way base J is in kinetoplastids, as the currently available *B. minutum* assembly is of too poor quality to allow such analysis.

What is the precise role of histones, DVNPs and HLPs in dinoflagellate genomes? Here, we demonstrate decreased chromatin accessibility over certain regions in the genome as well as the existence of nucleosome-like structures, which suggests the presence of nucleosomes along the genome. However, mapping of histone/DVNP/HLP occupancy using chromatin immunoprecipitation (ChIP) will be needed to generate direct genome-wide profiles of the distribution of these proteins along the genome. Currently this is precluded due to the extreme sequence divergence of dinoflagellate histone proteins ^29^, which makes existing anti-histone antibodies unreliable reagents for carrying out such experiments in dinoflagellates. Establishing the absolute levels of protection/occupancy in dinoflagellate genomes, through the application of methylation-based (especially long read-based) and enzymatic approaches ^99^ will also be highly valuable.

## Methods

### *B. minutum* cell culture

The clonal axenic *Symbiodinium*/*Breviolum minutum* strain SSB01 was used in all experiments. Stock cultures were grown as previously described ^100,101^ in Daigo’s IMK medium for marine microalgae (Wako Pure Chemicals) supplemented with casein hydrolysate (IMK+Cas) at 27 ^*°*^C at a light intensity of 10 *μ*mol photons m^*−*2^ s^*−*1^ from Philips ALTO II 25-W bulbs on a 12-h-light:12-h-dark cycle. The medium was prepared in artificial seawater (ASW).

### Genomic DNA isolation

*B. minutum* genomic DNA was isolated as previously described ^100^. Briefly, cells were centrifuged at 1,000 *g* for 5 minutes, then resuspended in 500 *μ*L 1*×* Cell Lysis Buffer (prepared by mixing equal volumes of 2*×*Cell Lysis Buffer – 2% SDS, 400 mM NaCl, 40 mM EDTA, 100 mM Tris-HCl, pH 8.0 – and H_2_O) and vortexed. The lysed cells were mixed with an equal 500 *μ*L volume Phenol:Chloroform:Isoamyl alcohol (25:24:1), and mixed well by inverting a few times. The phases were centrifugation at 13,000 *g* for 5 minutes, then the top phase was transferred to a new tube and treated with 4 *μ*L Ribonuclease A (20 mg/mL) by incubating for 30 minutes at 37 ^*°*^C.

DNA was purified by adding an equal volume Phenol:Chloroform:Isoamyl alcohol (25:24:1), mixing well and centrifuging at 13,000 *g* for 5 minutes, then transferring the top layer to a new tube, to which Phenol:Chloroform:Isoamyl alcohol (25:24:1) was added again, and the centrifugation and top phase isolation was repeated. Then 2.5*×*volumes of 100% EtOH were added and the mixture was incubated on ice for 30 minutes or at -20 ^*°*^C overnight. The solution was then centrifuged at 13,000 *g* at room temperature for 20 minutes, the pellet was washed with 70% EtOH, dried on air and resuspended in 50 *μ*L H_2_O.

### ATAC-seq experiments

ATAC-seq experiments were performed following the omni-ATAC protocol ^81^.

Briefly, *∼*100K *B. minutum* cells were centrifuged at 1,000 *g*, then resuspended in 500 *μ*L 1*×*PBS and centrifuged again. Cells were then resuspended in 50 *μ*L ATAC-RSB-Lysis buffer (10 mM Tris-HCl pH 7.4, 10 mM NaCl, 3 mM MgCl_2_, 0.1% IGEPAL CA-630, 0.1% Tween-20, 0.01% Digitonin) and incubated on ice for 3 minutes. Subsequently 1 mL ATAC-RSB-Wash buffer (10 mM Tris-HCl pH 7.4, 10 mM NaCl, 3 mM MgCl_2_, 0.1% Tween-20, 0.01% Digitonin) were added, the tubes were inverted several times, and nuclei were centrifuged at 500 *g* for 5 min at 4 ^*°*^C.

Transposition was carried out by resuspending nuclei in a mix of 25 *μ*L 2*×* TD buffer (20 mM Tris-HCl pH 7.6, 10 mM MgCl_2_, 20% Dimethyl Formamide), 2.5 *μ*L transposase (custom produced) and 22.5 *μ*L nuclease-free H_2_O, and incubating at 37 ^*°*^C for 30 min in a Thermomixer at 1000 RPM.

Transposed DNA was isolated using the MinElute PCR Purification Kit (Qiagen Cat# 28004/28006), and PCR amplified as previously described ^81^. Libraries were purified using the MinElute kit, then sequenced on a Illumina NextSeq 550 instrument as 2*×*36mers or as 2*×*75mers.

### ATAC-seq control experiments

Genomic DNA controls for ATAC-seq were generated by transposing purified gDNA. Briefly, 100 ng of gDNA were mixed with 2 *μ*L Tn5, 25 *μ*L 2*×*TD buffer and H_2_O for a total volume of 50 *μ*L, then incubated at 55 ^*°*^C for 5 minutes. The reaction was stopped by immediately proceeding with DNA isolation using the MinElute kit. Libraries were generated as described above for ATAC-seq.

### Genome assemblies

Datasets were processed against either the original *B. minutum* assembly ^35^ or against the Hi-C scaffolded assembly for *B. minutum* previously described ^40^, which is based on the original fragmented assembly for this species ^35^ and scaffolded into chromosome-level contigs using Hi-C data following established protocols ^102^.

### General analysis procedures

Browser tracks generation, fragment length estimation, and other analyses were carried out using custom-written Python scripts (https://github.com/georgimarinov/GeorgiScripts).

### Mappability track generation

Mappability tracks were generated as by tiling the whole genome with reads of length *RL* starting at each position. These reads were the mapped back to the genome using the same settings used for processing real datasets. Average mappability over each position was calculated as the ratio *RC/RL* between its read coverage *RC* and the read length *RL*.

ATAC-seq data processing

Demultipexed FASTQ files were mapped as 2*×*36mers using Bowtie ^103^ with the following settings: `-v 2 -k 2 -m 1--best --strata -X 1000`. Duplicate reads were removed using `picard-tools` (version 1.99). This mapping generated a set of uniquely mapping alignments only.

For the purpose of the analysis of multimappers, alignments were generated with unlimited alignment multiplicity with the following settings: `-v 2 -a --best --strata-X 1000`.

Normalization of multimappers was performed using the previously described ^104,105^ method of dividing each alignment by its read multiplicity, i.e:

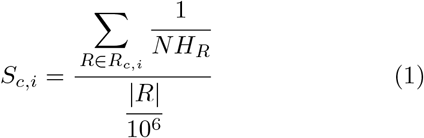

Where *S*_*c,i*_ is the signal score for position *i* on chromosome *c* (in RPM, or Read Per Million mapped reads units),|*R*| is the total number of mapped reads, |*R*_*c,i*_| is thenumber of reads covering position *i* on chromosome *c*, and *NH*_*R*_ is the number of locations in the genome a read maps to.

### ATAC-seq peak calling

Peak calling was carried out using MACS2^106^, with the gDNA library as a control, and with the following settings:-g 569785352 -f BAMPE. Differentially accessible regions were identified using DESeq2^107^.

### Analysis of positioned nucleosomes

The analysis of positioned nucleosomes was carried out using NucleoATAC ^93^. We used the low resolution nucleosome calling program nucleoatac occ with default parameters that requires ATAC-seq data and genomic windows of interest, and returns a list of nucleosome positions based on the distribution of ATAC-seq fragment lengths centered at these positions. Sliding windows of 1 kbp in steps of 500 bp were taken as inputs, and redundant nucleosome positions were eventually discarded. V-plots were made by aggregating unique-mapping ATAC-seq reads centered around the positioned nucleosomes, and mapping the density of fragment sizes versus fragment center locations relative to the positioned nucleosomes as previously described ^93,94^.

### MeDIP-seq experiments

To prepare inputs for MeDIP-seq experiments, gDNA was first sonicated using a Qsonica S-4000 with a 1/16” tip for 3 minutes, with 10 second pulses at intensity 3.5, and 20 seconds rest between pulses. The IP procedure was adapted from the protocol for ChIP-seq as previously described ^108^.

For each reaction, 100 *μ*L of Protein A Dynabeads (ThermoFisher Cat # 10002D) were washed 3 times with a 5 mg/mL BSA solution. Beads were then resuspended in 1 mL BSA solution and 5 *μ*L of *α*-5-hmU antibody (Abcam Cat # ab19735) were added. Coupling of antibodies to beads was carried out overnight on a rotator at 4 ^*°*^C. Beads were again washed 3 times with BSA solution and resuspended in 100 *μ*L of BSA solution.

Sheared genomic DNA (*∼*1 *μ*g 1:1 mix of *B. minutum* and *Homo sapiens*) was end repaired and adapters were ligated to it following the procedure of the NEBNext Ultra II DNA Library Prep Kit for Illumina (NEB, E7645S), purified using AMPure XP beads and eluted in 50 *μ*L of H_2_O, and then denatured at 98 ^*°*^C for 10 minutes. DNA was then immediately placed on ice, resuspended in 850 *μ*L RIPA buffer (1*×*PBS, 1% IGEPAL, 0.5% Sodium Deoxycholate, 0.1% SDS, Roche Protease Inhibitor Cocktail) and added to the beads, then incubated overnight on a rotator at 4 ^*°*^C.

Beads were washed 5 times with LiCl buffer (10 mM Tris-HCl pH 7.5, 500 mM LiCl, 1% NP-40/IGEPAL, 0.5% Sodium Deoxycholate) by incubating for 10 minutes at 4 ^*°*^C on a rotator, then rinsed once with 1*×*TE buffer. Beads were then resuspended in 200 *μ*L IP Elution Buffer (1% SDS, 0.1 M NaHCO_3_) and incubated at 65 ^*°*^C in a Thermomixer (Eppendorf) with interval mixing to dissociate antibodies. Beads were separated from the DNA solution by centrifugation, and DNA was purified using the MinElute kit.

Library generation was completed by carrying out PCR following the rest of the steps of the NEBNext Ultra II DNA Library Prep Kit protocol, using 15 cycles of amplification. Final libraries were purified using AMPure XP beads.

Several control libraries were prepared – “Input” from the gDNA that was used as input to the immunoprecipitation, “Depleted” from the supernatant from the first bead separation after the incubation of DNA with beads, and “IgG”, generated from a parallel immunoprecipitation reaction that used only Protein A beads (without a primary antibody)

### MeDIP-seq data processing

MeDIP-seq libraries processing was carried out in the same way as that of ATAC-seq datasets.

### 5-hmU chemical mapping experiments

Chemical mapping of 5-hmU as carried out following the previously described by Kawasaki et al. chemical conversion method ^98^. Briefly, sheared genomic DNA was used as input and end prep and adapter ligation were carried out using the NEBNext Ultra II DNA Library Prep Kit. After the ligation step, DNA was purified using AMPure XP beads and eluted in 50 *μ*L of H_2_O. DNA denaturation was performed by adding NaOH to a final concentration of 0.05M and incubating at 37 ^*°*^C for 30 minutes. Oxidation was carried out by adding 2 *μ*L of KRuO_4_ solution (15 mM in 0.05 M NaOH) and incubating for 30 minutes at room temperature. Oxidized DNA was purified using AMPure XP beads and extension was carried out by mixing 13.5 *μ*L DNA, 1.6 *μ*L 100 mM MgSO_4_, 2 *μ*L NEB Index Primer, 2 *μ*L 10*×*ThermoPol Reaction Buffer (NEB), 0.5 *μ*L 10 mM dNTP mix, and 0.4 *μ*L Bst DNA Polymerase, Large Fragment (NEB), then incubating for 1 hour at 37 ^*°*^C. PCR amplification was carried out using the NEB Ultra DNA Library Prep Kit, with 12 cycles of PCR. Final libraries were purified using AMPure XP beads.

### Processing of 5-hmU chemical mapping datasets

The slamdunk package ^109^ (https://t-neumann.github.io/slamdunk/), which was originally developed for the analysis of SLAM-seq ^110^ datasets (the SLAM-seq protocol also generates T*→*C conversions) was adapted to estimate 5-hmU conversion levels.

First, the genome was tiled into 500-bp bins starting every 100 bp. Second, sequencing reads were trimmed of adaptors using Trim Galore, and used as input to slamdunk together with the genome tiling with the following settings: `--max-read-length 75 -5 9 -n 1000000 -m --skip-sam`.

### Repeat annotation

Repeats were identified de novo from the scaffolded assembly using RepeatModeler-2.0.1 with default parameters. Repeat annotations were subsequently generated using RepeatMasker-4.1.1^111^ with RMBlast-2.10.0 as the sequence search engine.

### Analysis of ATAC-seq and MeDIP-seq data in repeat space

Sequencing datasets were analyzed in repeat space as previously described ^105^. Briefly, reads were mapped to consensus repeat sequences with relaxed settings (-e 200 instead of-v 2) and with unlimited multimappers. Normalization of multimapping reads was carried out as above.

### Heterologous expression of DVNPs in yeast

The MS46 *S. cerevisiae* strain ^112^ was used for all experiments.

For the experiments in Supplementary Figure 4, each DVNP was expressed from a SIVu-WTC846::TetPr-DVNP-3xNLS-linker-3PK construct that was integrated in a single copy at the URA locus of the MS46 stain. The WTC846::TetPr promoter is reported previously ^113^. Cells were grown in YPD media at 30 ^*°*^C overnight and expression was induced by the addition of 200 nM anhydrotetracycline. Cells were collected 5 hours after induction. Untransformed MS46 cells were used as control.

For the experiments in Supplementary Figure 5, each DVNP was expressed from a multicopy pRS416-GAL1pr-DVNP-3xHA-NLS plasmid, as first reported in by Irwin et al. ^83^, that was transformed into MS46. Cells were grown in synthetic media lacking uracil + 2% raffinose at 30 ^*°*^C and cultured overnight before expression was induced by the addition of 2% galactose to the media. Cells were collected 7 hours after induction. MS46 transformed with an empty pRS416-GAL1pr construct was used as control.

### Yeast SMF experiments

Yeast SMF experiments were carried out as previously described 96,114–116

A 1:1 mixture of *S. cerevisiae* cells expression DVNPs and *Candida glabrata* cells (used as a control for normalization, as previously described ^86^) amounting to a total of 2.5 *×* 10^8^ cells was used as input. Cells in log phase (OD_660_*≤*1.0) were first centrifuged at 13,000 rpm for 1 minute, then washed with 100 *μ*L Sorbitol Buffer(1.4 M Sorbitol, 40 mM HEPES-KOH pH 7.5, 0.5 mM MgCl_2_), and centrifuged again at 13,000 rpm for 1 minute. Cells were then spheroplasted by resuspending in 200 *μ*L Sorbitol Buffer with DTT added at a final concentration of 10 mM and 0.5 mg/mL 100T Zymolase, followed by incubating for 5 minutes at 30 ^*°*^C at 300 rpm in a Thermomixer. The pellet was centrifuged for 2 minutes at 5,000 rpm, washed in 100 *μ*L Sorbitol Buffer, and centrifuged again at 5,000 rpm for 2 minutes.

Cells were then resuspended in 100 *μ*L ice-cold Nuclei Lysis Buffer (10 mM Tris pH 7.4, 10 mM NaCl, 3 mM MgCl_2_, 0.1 mM EDTA, 0.5% NP-40) and incubated on ice for 10 minutes. Nuclei were then centrifuged at 5000 rpm for 5 min at 4 ^*°*^C, resuspended in 100 *μ*L cold Nuclei Wash Buffer (10 mM Tris pH 7.4, 10 mM NaCl, 3 mM MgCl_2_, 0.1 mM EDTA), and centrifuged again at 5,000 rpm for 5 min at 4 ^*°*^C. Finally, nuclei were resuspended in 100 *μ*L M.CviPI Reaction Buffer (50 mM Tris-HCl pH 8.5, 50 mM NaCl, 10 mM DTT).

Nuclei were then first treated with M.CviPI (GpC methyltransferase) by adding 200 U of M.CviPI (NEB), SAM at 0.6 mM and sucrose at 300 mM, and incubating at 30 ^*°*^C for 7.5 min. After this incubation, 128 pmol SAM and another 100 U of enzymes were added, and a further incubation at 30 ^*°*^C for 7.5 min was carried out. Immediately after, M.SssI treatment (CpG methyltransferase) followed, by adding 60 U of M.SssI (NEB), 128 pmol SAM, MgCl_2_ at 10 mM and incubation at 30 ^*°*^C for 7.5 min.

The reaction was stopped by adding an equal volume of Stop Buffer (20 mM Tris-HCl pH 8.5, 600 mM NaCl, 1% SDS, 10 mM EDTA).

HMW DNA was isolated using the MagAttract HMW DNA Kit (Qiagen; cat # 67563) following the manufacturer’s instructions.

Enzymatically labeled DNA was then sheared on a Covaris E220 and converted into sequencing libraries following the EM-seq protocol, using the NEBNext Enzymatic Methyl-seq Kit (NEB, Cat # E7120L).

### Yeast SMF data processing

Adapters were trimmed from reads using Trimmomatic ^117^ (version 0.36). Trimmed reads were aligned against a combined *S. cerevisiae sacCer3* plus *Candida glabrata* C_glabrata_CBS138 genome index using bwa-meth with default settings. Duplicate reads were removed using picard-tools (version 1.99). Methylation calls were extracted using MethylDackel (https://github.com/dpryan79/MethylDackel). Additional analyses were carried out using custom-written Python scripts (https://github.com/georgimarinov/GeorgiScripts).

Chemically mapped nucleosome positions in *S. cerevisiae* were obtained from Brogaard et al. 2012^118^ as previously described ^96^.

### Yeast ATAC-seq experiments

Yeast ATAC-seq experiments were carried out as previously described ^96,119^.

Briefly, ATAC-seq was carried out on the same nuclei isolated for SMF as described above (before resuspension in M.CviPI Reaction Buffer), by resuspending nuclei with 25 *μ*L 2*×*TD buffer (20 mM Tris-HCl pH 7.6, 10 mM MgCl_2_, 20% Dimethyl Formamide), 2.5 *μ*L transposase (custom produced) and 22.5 *μ*L nuclease-free H_2_O, and incubating at 37 ^*°*^C for 30 min in a Thermomixer at 1000 RPM. Transposed DNA was isolated using the DNA Clean & Concentrator Kit (Zymo, cat # D4014) and PCR amplified as described before ^81^. Libraries were then sequenced on a Illumina NextSeq instrument as 2*×*36mers or as 2*×*75mers.

## ATAC-seq data processing

FASTQ files were mapped against a combined *S. cerevisiae sacCer3* plus *Candida glabrata* `C_glabrata_CBS138` genome index as 2*×*36mers using Bowtie ^103^ with the following settings: `-v 2 -k 2 -m 1 --best --strata`. Duplicate reads were removed using `picard-tools` (version 1.99). Additional analysis was carried out as previously described ^120^.

## Author contributions

G.K.M. conceptualized the study, and carried out ATAC-seq, MeDIP and 5-hmU chemical mapping experiments. X.C. analyzed data. M.P.S. carried yeast DVNP expression experiments. T.X. carried out cell culture and DNA isolation. A.R.G, A.K. and W.J.G. supervised the study.

G.K.M. and X.C. wrote the manuscript with input from all authors.

## Acknowledgements

This work was supported by NIH grants (P50HG007735, RO1 HG008140, U19AI057266 and UM1HG009442 to W.J.G., 1UM1HG009436 to W.J.G. and A.K., 1DP2OD022870-01 and 1U01HG009431 to A.K.), the Rita Allen Foundation (to W.J.G.), the Baxter Foundation Faculty Scholar Grant, and the Human Frontiers Science Program grant RGY006S (to W.J.G). W.J.G. is a Chan Zuckerberg Biohub investigator and acknowledges grants 2017-174468 and 2018-182817 from the Chan Zuckerberg Initiative. Fellowship support provided by the Stanford School of Medicine Dean’s Fellowship (G.K.M.). This work is also supported by NSF-IOS EDGE Award 1645164 to A.R.G. and Carnegie Venture grant 10907 (to T.X. and G.K.M.).

The authors would like to thank Alexandro Trevino for supplying the *α*-5-hmU antibody, Nicholas Irwin and Patrick Keeling for providing the construct for expressing *Hematodinium* DVNP.5, as well as members of the Green-leaf, Kundaje, Pringle and Grossman laboratories for helpful discussion and suggestions regarding this work.

## Data Availability

Data associated with this manuscript have been submitted to GEO under accession number GSE241969.

## Code Availability

Custom code used to process the data is available at https://github.com/georgimarinov/GeorgiScripts.

## Supplementary Materials

### Supplementary Figures

**Supplementary Figure 1:**
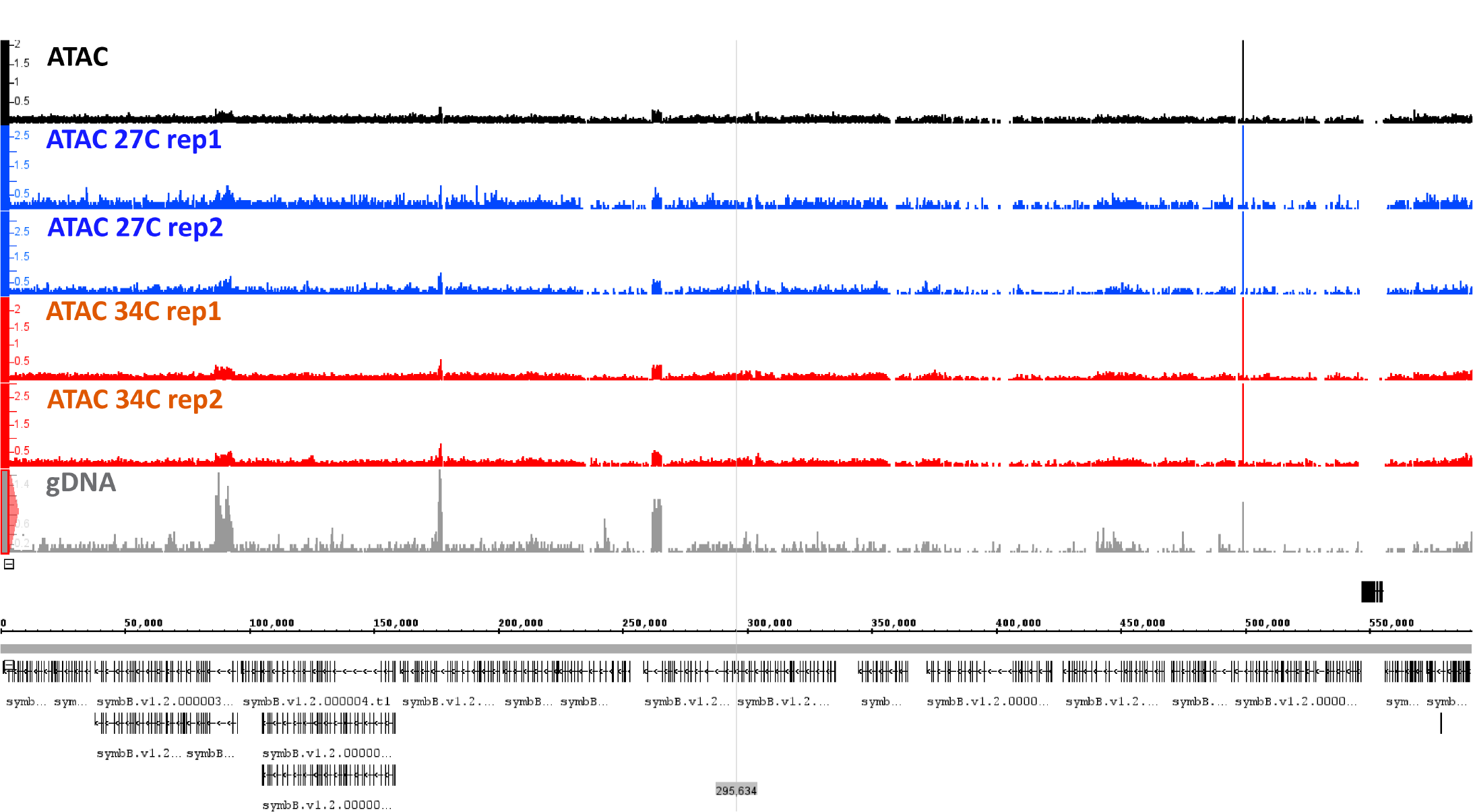
Representative genome browser view of ATAC-seq and gDNA control signal in the *B. minutum* genome.

**Supplementary Figure 2:**
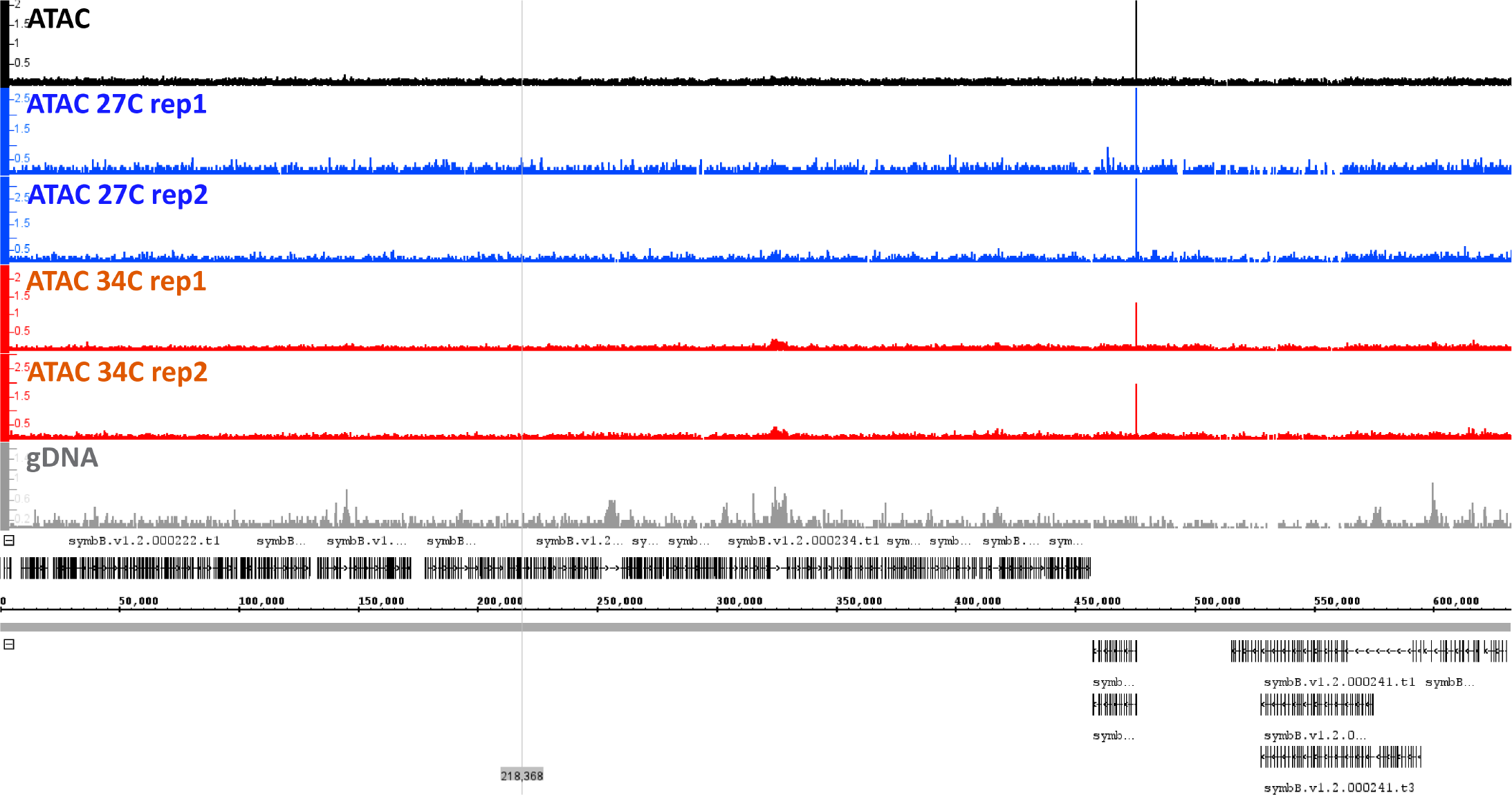
Representative genome browser view of ATAC-seq and gDNA control signal in the *B. minutum* genome.

**Supplementary Figure 3:**
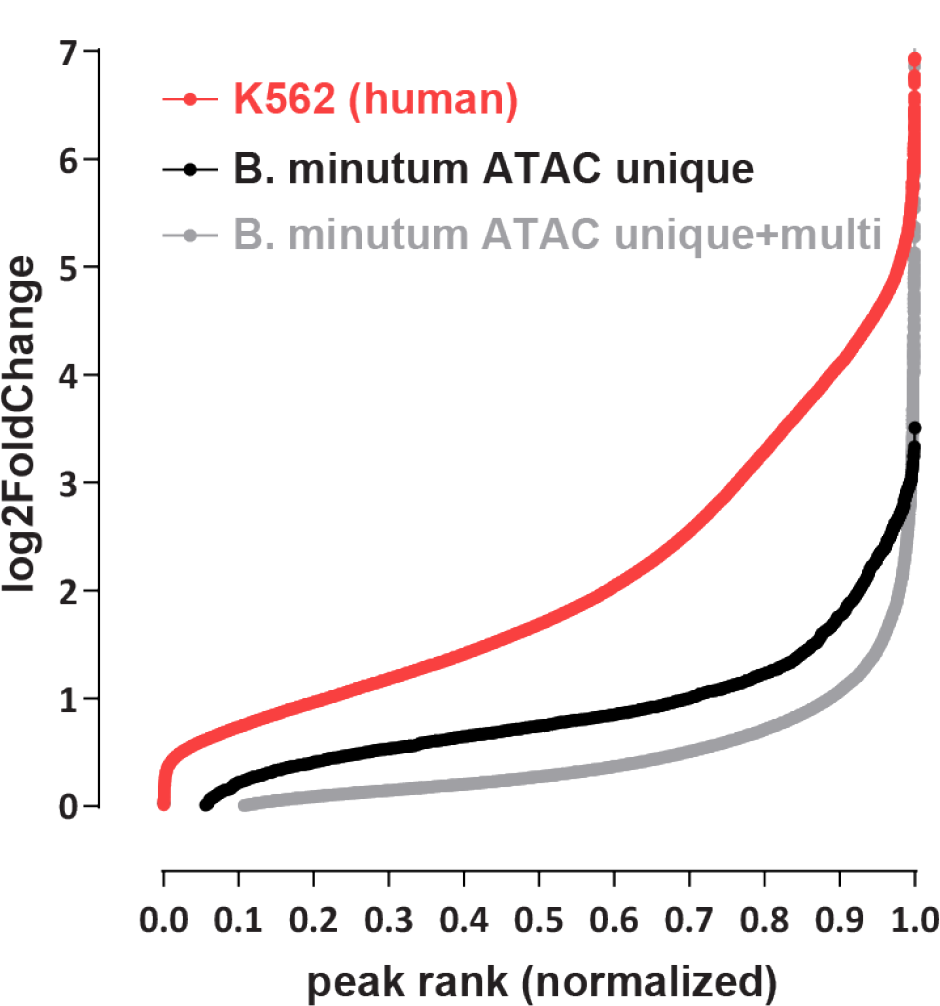
Relative degree of ATAC-seq enrichment in *B. minutum* versus a representative mammalian genome sample. Shown is the *log*_2_(fold change) ratio of ATAC-seq signal versus a negative control for a representative human ATAC-seq sample (K562 cell line from the ENCODE Project Consortium ^82^; dataset ID ENCFF512VEZ was used for ATAC and dataset ID ENCFF285UKJ – a whole genome bisulfite sequencing library – as a negative control, over peaks from dataset ID ENCFF695IGF). A separately sequenced gDNA control was generated for the *B. minutum* ATAC.

**Supplementary Figure 4:**
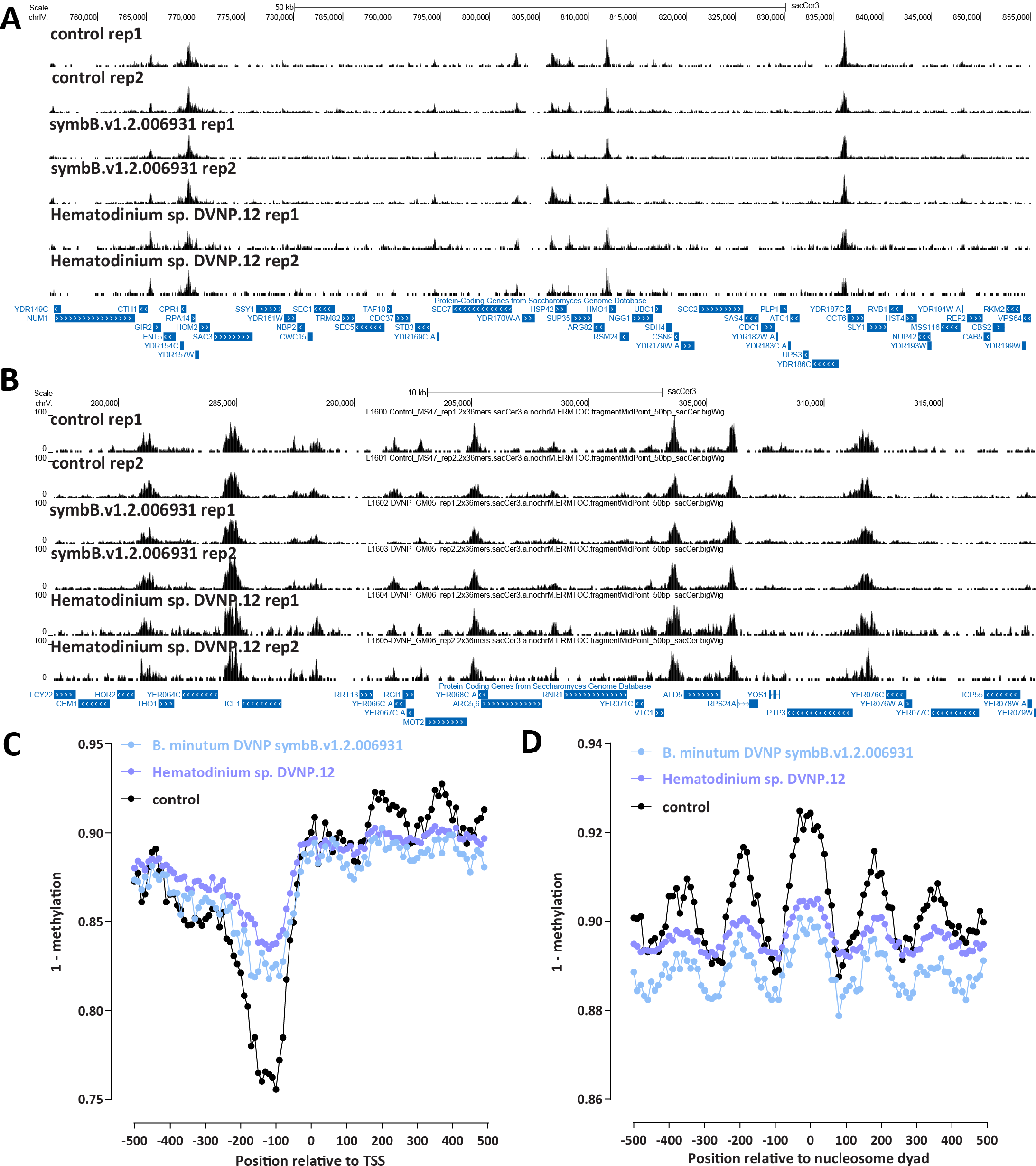
Effects of exogenous expression dinoflagellate DVNPs on chromatin accessibility in the yeast *S. cerevisiae*. (A-B) ATAC-seq profiles of *S. cerevisiae* expressing *B. minutum* DVNP symbB.v1.2.006931 and *Hematodinium* sp. DVNP.12 and control samples. (C) SMF profiles (corrected using average SMF methylation from the *Candida* internal control) over *S. cerevisiae* TSSs in *S. cerevisiae* expressing *B. minutum* DVNP symbB.v1.2.006931 and *Hematodinium* sp. DVNP.12 and control samples. (D) SMF profiles (corrected using average SMF methylation from the *Candida* internal control) over positioned *S. cerevisiae* nucleosomes in *S. cerevisiae* expressing *B. minutum* DVNP symbB.v1.2.006931 and *Hematodinium* sp. DVNP.12 and control samples.

**Supplementary Figure 5:**
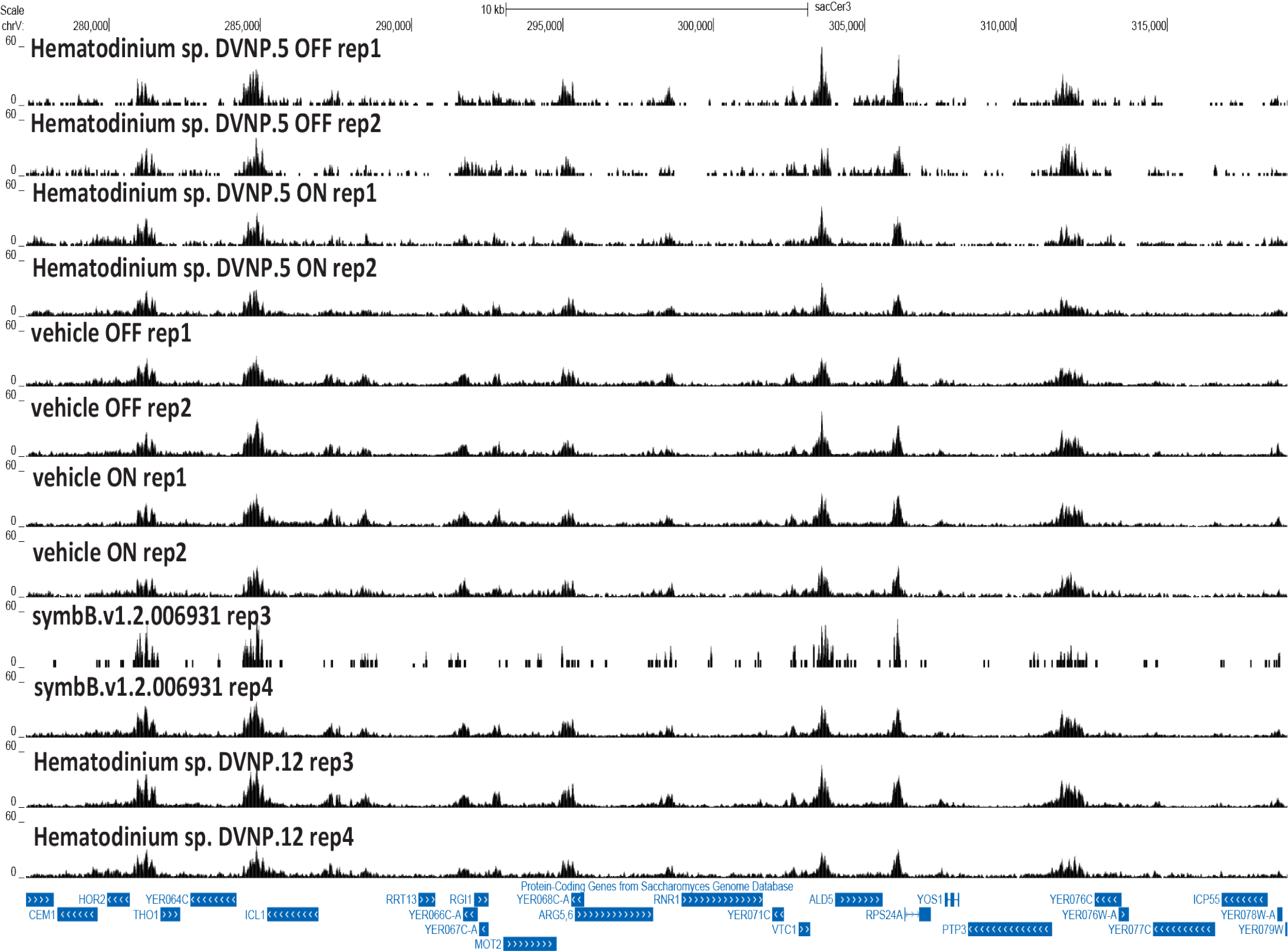
Effects of exogenous expression of dinoflagellate DVNPs on chromatin accessibility in the yeast *S. cerevisiae*. ATAC-seq profiles of *S. cerevisiae* expressing *Hematodinium* sp. DVNP.5 (from Irwin et al. 2018^83^) and a vehicle control, as well as additional replicates for *B. minutum* DVNP symbB.v1.2.006931 and *Hematodinium* sp. DVNP.12 and control samples. “OFF” and “ON” refer to cells in which the expression of *Hematodinium* sp.DVNP.5 is induced or not.

**Supplementary Figure 6:**
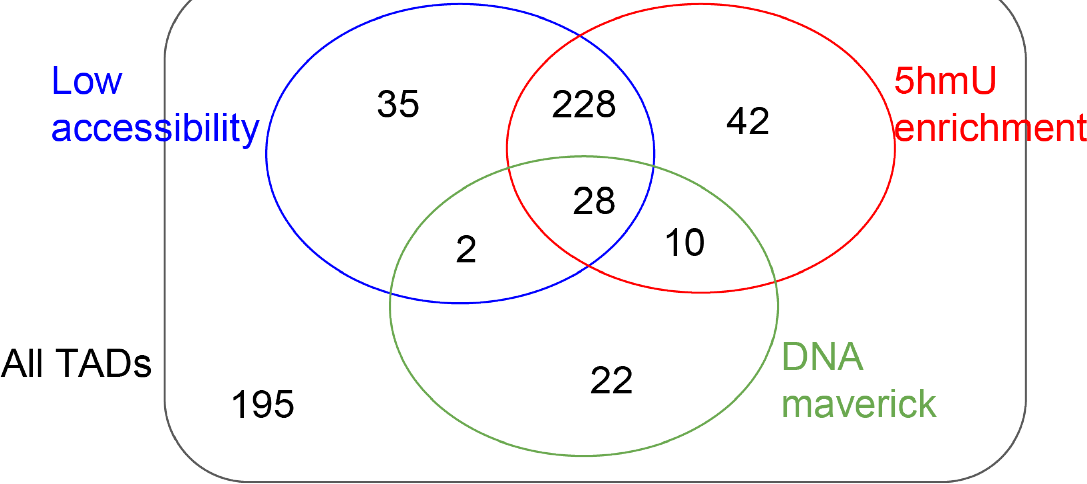
Overlap between dinoTAD boundaries, regions of low accessibility, and regions of high 5-hmU.

**Supplementary Figure 7:**
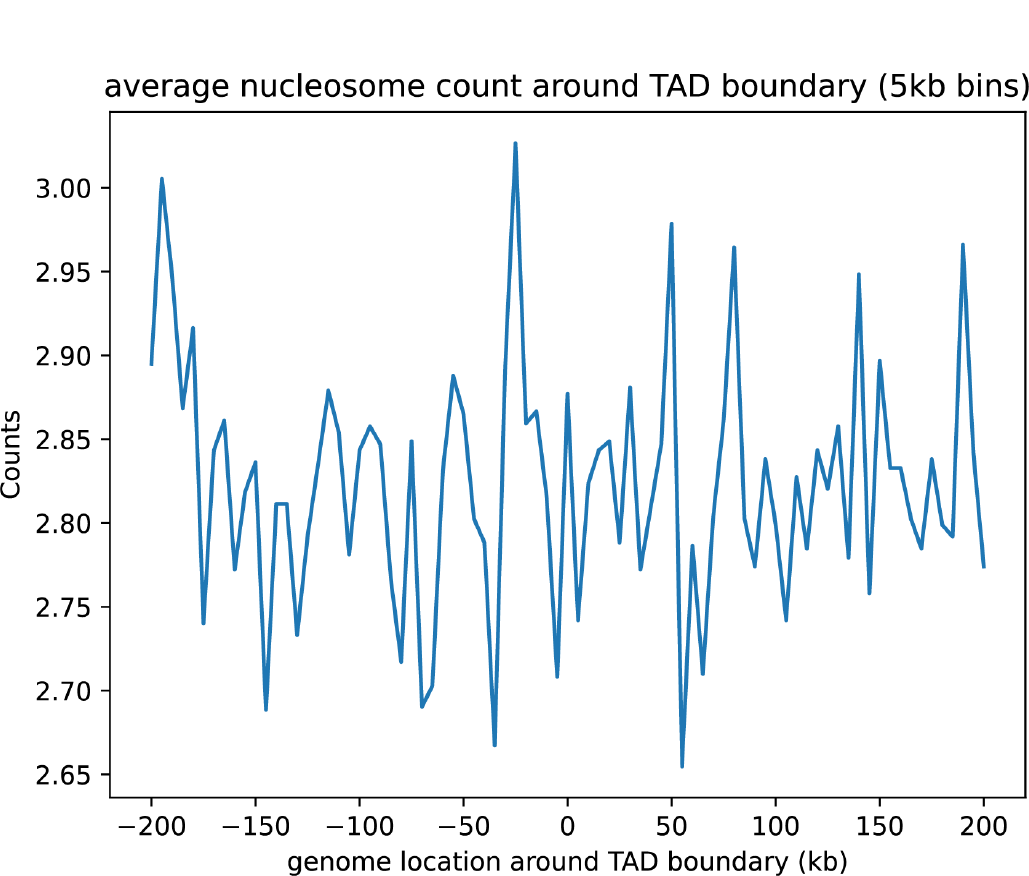
Positioned nucleosomes as a whole are not strongly enriched around dinoTAD boundaries.

**Supplementary Figure 8:**
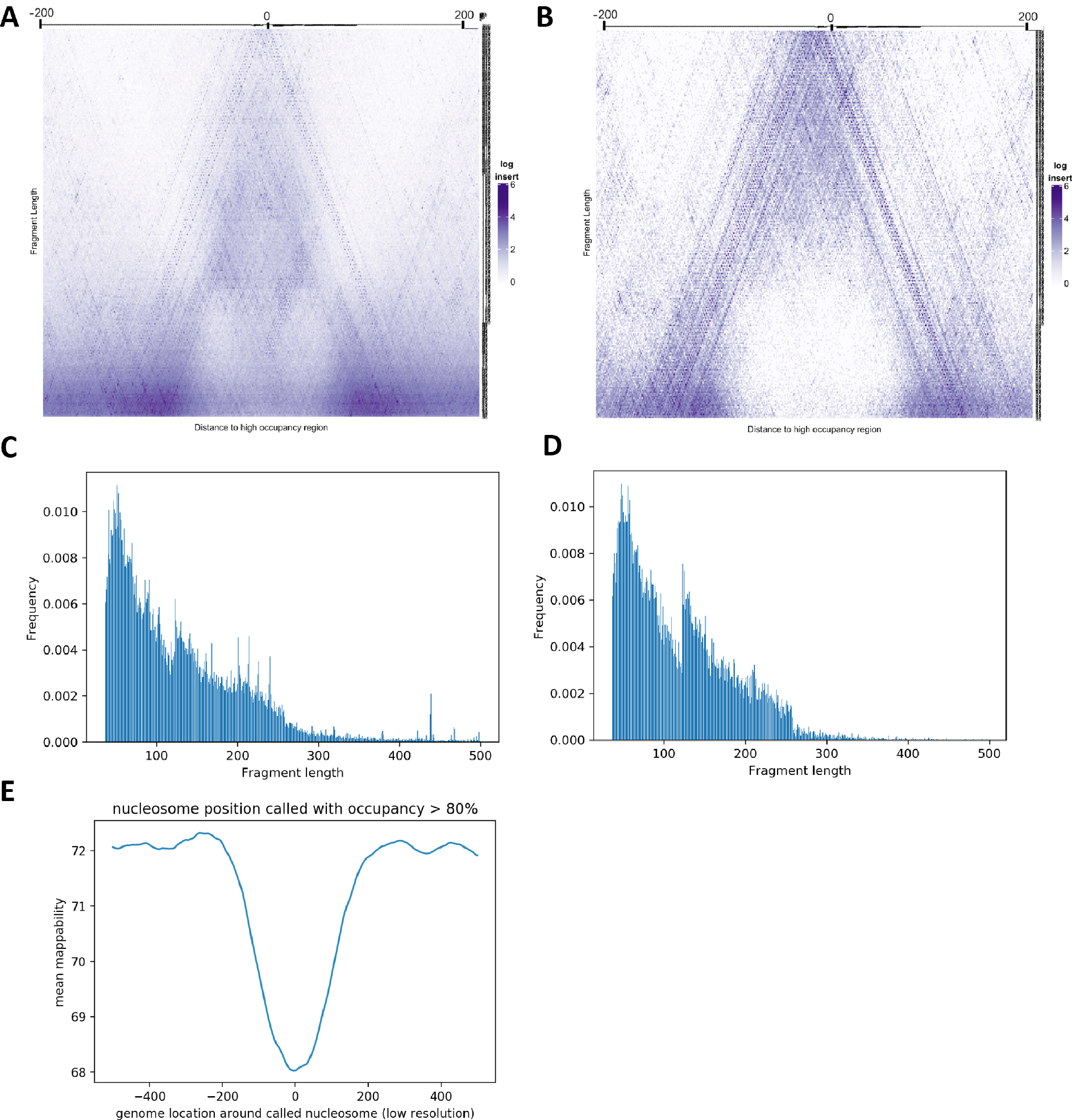
Properties of putative positioned nucleosomes in the *B. minutum* genome. (A) V-plot of low-resolution positioned nucleosomes (*n*=30,107) (B) V-plot of low-resolution positioned nucleosomes with minimum occupancy cutoff of 0.8 (*n*=2,166) (C) Fragment distribution over low-resolution positioned nucleosomes (D) Fragment distribution over high-resolution positioned nucleosomes with minimum occupancy cutoff of 0.8 (E) Average mappability (for reads of length 75 bp) over positioned nucleosomes.

